# Bypassing juvenility, first report of functional flowering and fruiting in six months old seedlings of *Cordia myxa L.(lasuda)*

**DOI:** 10.1101/2020.06.12.148122

**Authors:** Vipan Guleria

**Author notes:** (DR.Y S Parmar University of Horticulture and Forestry, Nauni (Solan)).

## Abstract

*Cordia myxa* L. is a medium-sized broad-leaved deciduous tree of flowering plant belongs to Boraginaceae. It grows naturally from dry desert India up to hills of Himalayas in India up to 1400 m elevation above mean sea level. Fruits are mostly used for pickle making and dried for its use in local off season vegetable recipes since long time. The ripe fruits are full of vitamins and its regular use is supposed to be helpful in good growth of hair. Lasuda preparations are, thus, good for people whose constitution might have tendency to go bald. In addition to fruit, Lasuda bark and roots are also very effective as a local remedy against cough, cold and various other ailments connected with indigestion and throat problems. At regional research laboratory of Dr. Y. S. Parmar University of Horticulture and Forestry, Nauni (India) we are working for the last 12 years on various aspects like propagation and development of promising strains of *Cordia myxa*. Grafting/budding techniques have been standardized to produce true to type precocious plants which bear flower and fruits in two to three years. However, flowering and fruiting has been observed in six months old seedling of seed origin, which can be ascribed to biochemical or cellular changes. Early flowering and fruiting is a rare phenomenon in tree seedlings of the species which otherwise flower at the age of 7-8 years. This could be very useful for manipulating the species at gene as well as physiological level in future to get early fruits and breeding of the species.

## Introduction

*Cordia myxa* is traditional underutilized fruit tree with multiple uses such as vegetable, fodder and soil binder tree of lower hill regions of Himachal Pradesh. In India, an average seedlings start bearing flowers at the age of around seven to eight years. Lasuda tree flowers during April– May. The inflorescence, mostly terminal, is, white in color. Individual florets are nearly 5 mm in diameter. At places these are somewhat hairy and white. Being a deciduous plant, the species bears male and female flowers on the same tree. Bole is tortuous or straight; bark grey, cracked; branches spreading, forming a dense crown; branchlets hairy, later glabrous, with very prominent leaf scars. Leaves alternate, simple; stipules absent; petiole 0.5–4.5 cm long; blade broadly ovate to orbicular, sometimes obovate, 3–18 cm × 3–20 cm, base rounded to cordate or cuneate, apex rounded to obtusely acuminate, margins entire to toothed, glabrous above, glabrous to velvety hairy below. Inflorescence a lax terminal or short lateral panicle, 3.0–8.5 cm long, many-flowered; bracts absent. Flowers unisexual, regular, white to creamy; pedicel 1–2 mm long; male flowers with campanulate calyx 4.5–5.5 mm long, 3-lobed, shortly hairy inside, glabrous outside, corolla tube 3.5–4.5 mm long, lobes 5, elliptical, c. 5 mm × 2 mm, reflexed, stamens inserted at corolla throat, exerted, filaments 1.5–3.5 mm long, ovary rudimentary; female flower with tubular-campanulate calyx 6– 8.5 mm long, irregularly 3–4 toothed, densely hairy inside, glabrous outside, corolla tube 4.5–6.5 mm long, lobes 4–6, elliptical to obovate, 5–7 mm long, reflexed and rolled up, staminodes with sterile anthers, ovary superior, ellipsoid to obovoid, 4-celled, style 8–9 mm long, with 4 stigmatic branches 4–5 mm long. Fruit a globular to ovoid drupe 2.0–3.3 cm long, apiculate, enclosed at base by the accrescent calyx, yellow, apricot or blackish when ripe, pulp almost transparent, mucilaginous, sweet-tasting (Troup,1921). Pyrene broadly ellipsoid to globose, c. 12 mm long, deeply wrinkled, 1–2-seeded. It is a delicacy on rural “Pahadi” cuisine in summer and rainy season in western Himalayas. Fruits are used for pickle making and dried for its use as an off season vegetable. It extends from Jammu, Himachal Pradesh to Uttrakhand and in dry part of Rajesthan of Indian Subcontinent. It is member of family Boraginaceae. It contains Stigma-sterol upto 5.86 per cent that Prevents certain types of cancers including ovarian, prostate, breast, and colon cancers, Possess certain antioxidant, hypoglycemic, hyperlipedimic, and thyroid inhibiting properties (Shwaish and Al-Imarah, 2017). *Cordia myxa* has certain analgesic, anti-inflammatory, immunomodulatory, antimicrobial, antiparasitic, insecticidal, cardiovascular, respiratory, gastrointestinal and protective effects(Al-Snafi, 2017). It playing an important role in the rural economy of arid regions (Peter,2007). The fruits can be easily dried after blanching for use during off season. it can give good returns even under rainfed marginal and less productive areas of hills. *Cordia dictoma*, Ehrenb. syn. *C. rothii* Roem. et Schult.; *Cordia macleodii* (Griff.) Hook., Thomson and *Cordia sebestena* L. are other important species within the genus *Cordia*., The importance of Lasuda fruit is well recognized both in rural as well as urban masses (Bhatnagar *et al*, 2017). Its fruit is edible, sweet and mucilaginous. The tender mature fruits are mostly pickled and also used for vegetable purpose. Fruits have medicinal feature and are considered as anthelmintic, diuretic, demulcent and expectorant (Oza and Kulkarni. 2016). They are rich source of carbohydrates, phosphorous, Sugar and calcium. Study and evaluation of the variation is the first step for any breeding programme and tree breeding depends on the existing variability in the nature under different edaphic-climatic conditions. The low hill Himalayan region is rich in its biodiversity under natural habitats where seedling trees come up in plethora naturally. Very scanty work on morphological variation in fruit quality and conservation has been reported on lasuda genotypes in this region.

## 2. Materials and Methods

Ripened fruits of *Cordia myxa* L were collected from wild growing trees in the month of June-July. The fruits were depulped and endocarps were separated and dried in open sun. endocarps are soaked in two per cent Bavistin solution for ten minutes were allowed to germinate in coco peat with germination trays. In total, 2,000 endocarps were used for germination studies. The Germination trays were kept in germination chamber on and monitored regularly. Germination started after 10-15 days in the month of September. The germinated endocarps were pricked in polybags of size 9”× 4” for further growth and development under protected conditions till the winter were over. The plants were observed regularly. Growth as well as development parameters viz., height, number of branches, spike length and number of flowers were recorded on species

## Results and Discussion

Around 1500 seedlings were kept under the similar growing conditions, none of the seedlings except one produced flowers as well as fruits at the age of six months i.e. in the first season of growing season after germination (**Figure 1**). The data recorded with regard to growth and develop-ment parameters of this unique seedling are presented in **Table 1**. The height of the seedling was just 13.9 cm whereas six numbers of branches were recorded. Nine number of flowers were counted and 4 number of fruits were developed (44.4% conversion rate). The fruit diameter in first week of May is around 10-13 mm. The seedlings has around 10 number of leaves in total. The species concerned is heavily destroyed by the insect attack and mass propagation is big problems due to poor regeneration apart from this the authors are working on the species for the last many years for development of improved cultivars and standardization of propagation techniques through seed and vegetative means. The grafted/ budded plant of Lasuda with higher productivity and nutritional value has been produced by the authors for quality and early fruiting as fruit is the main part of the plant from medicinal point of view. In present case, the seedling plant of Lasuda produced flowers at the age of just six months or in its first year which is a unique phenomenon. This is the first ever reported seedling plant of *Cordia myxa* came in flowering at such a young age other-wise plants of same origin bear flowers after 7-8 years of age. Over the years, more than 10,000 Lasuda plants have been produced by the authors from seeds yet it is the first instance when flowering has been observed in just six months old seedling

**Table 1.** Age, growth and development parameters of Harar seedling Parameter.

| **Table 1. Age, growth and development parameters of Harar seedling.** | **Value** |
| --- | --- |
| **Parameter** |  |
| Age of plant | 6 months |
| Height | 13,9 cm |
| Number of branches | 06 |
| Number of flowers | 9 |
| Number of fruits | 4 |
| Fruit diameter ( first week of May) | 11 |
| Number of leaves on seedlings | 10 |
| Fruit diameters (mm) | 10-13 |

**Figure 1:**
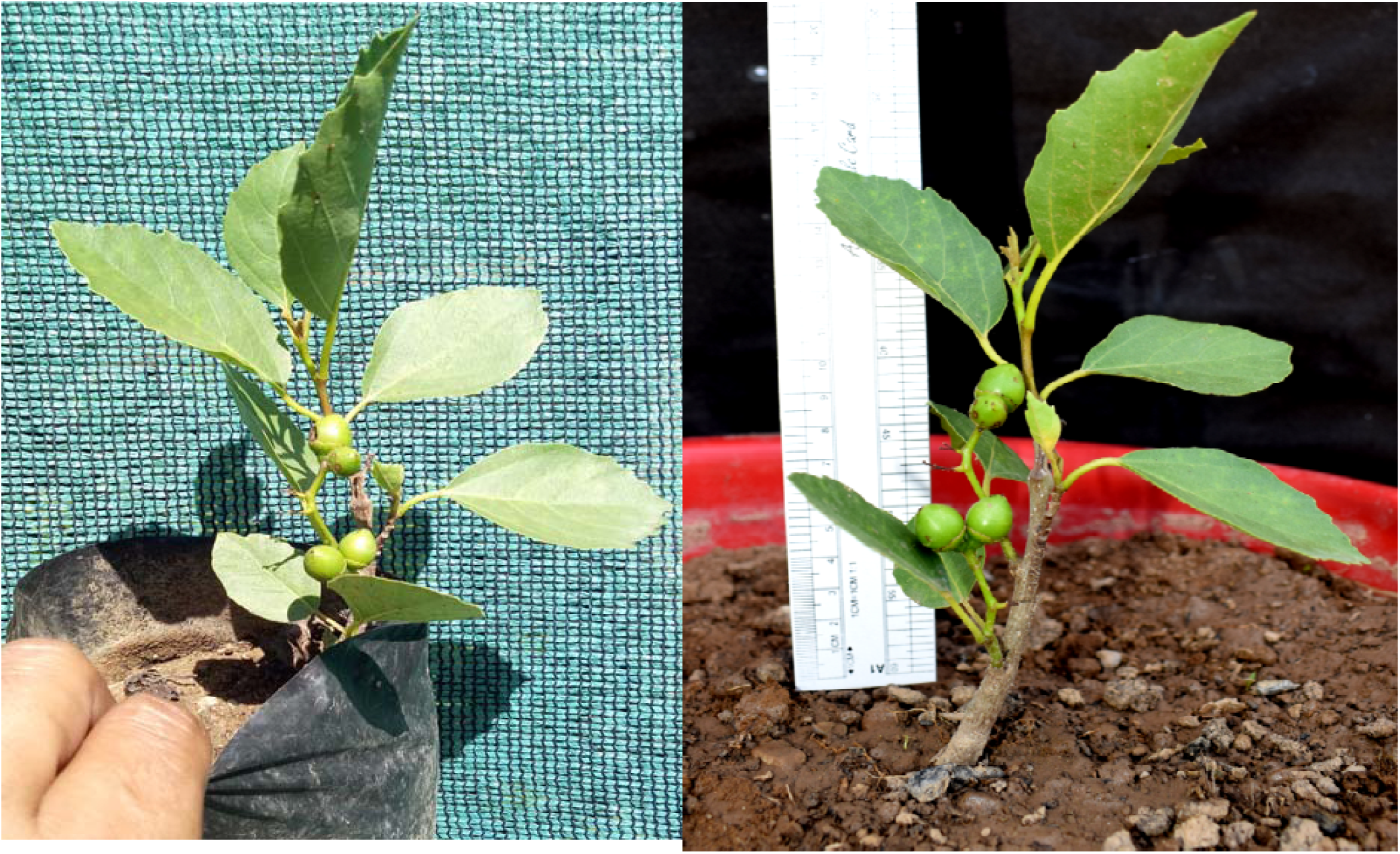
Flowering in six months old *Cordia myxa* seedling.

After certain age i.e. after some years each species of higher plants enter a period of reproductive growth where, leaf bud tissue changes its physiological state to become flower bud tissue, and then develops into a floral organ. The biochemical and cellular changes such as we know that florigen level decide the initiation of flowering in trees, this process is called flower bud differentiation (Guo et al., 2015). Flower bud differentiation is a complex process of morphogenesis. It is triggered by various factors, such as photoperiod, vernalization, nutrition, and water status, and is accomplished by the interaction and coordination of hormones and PAs (Xu, 2015). The floral primordial form and develop at the shoot apex as a result of bio-chemical and cellular changes leading to the production of the floral organs. Recent studies clearly indicate that the functions of plant hormones are not restricted to a particular stage, and a complex network of more than one plant hormone viz. Carbohydrates, Gibberellins and Cytokinins etc. are involved in controlling various aspects of fruit (Kumar, Khurana, and Sharma. (2014). The flowering physiology and biology of *Picea abies* has been found affected by environmental factors like temperature and rainfall by *Tiren (1935)*. Pharis and King (1985) concluded that the Cone formation can be promoted in juvenile plants of several gymnosperm families by application of gibbrrelic acids.. Hilton(2008) achieved over 90% flowering by applying the calcium carbide to 12-month-old pineapple plants one month before normal flowering. Bernier *et al* (1997) reported that the Cytokinins are completely unable to induce flowering, it appears that there is a multi-component floral stimulus in *Sinapis alba*. The floral hormone or the floral stimulus causes or evokes flowering and which is common to all higher plants(Lang A. 1965). Only Flowering in pine (conifer spp) and *Terminalia chebula* (broad leave spp) plant of less than one year of age has been reported by Johnson and Critchfield (1978) and Sharma *et al* (2012). Citrus plants recorded flowering in its first year but failed to fruit and hoped that the further investigations will help in solving the slow citrus breeding (Furr, Cooper and Reece,1947). Single major gene for precocious flowering in *Dalbergia sissoo* has been reported (Vakshasya and Rawat 1986). Many studies have shown that exogenous PAs and PA synthesis inhibitors can affect flower bud differentiation. Exogenous PAs were shown to accelerate the process of flower bud differentiation, and high PA contents in apical buds were beneficial for the initiation and maintenance of flower bud differentiation in *Chrysanthemum* (Xu, 2015). Cecich, Kang and Chalupka (1994) reported that flower initiation marks the transition from vegetative to reproductive growth in seed plants. It is thus a crucial event in the life of these plants, particularly so because of the peculiar relation of vegetative and reproductive development in seed plants which is in turn an outcome of the morphological nature of the flower.

The longer time in onset of flowering in trees species pose a substantial obstacle to research and breeding in time bound frame Zhang et al (2010). Cecich (1981) has reorted that the development of self breeding lines is a key step in crop improvement programmes because it allows the production of true breeding varieties and reproducible hybrids. Flowering is heritable character as reported by Johnson and Critchfield (1978) and if we go by it then the present plant type good and rare opportunity to genetic studies to fix the gene responsible for flowering and manipulations can bring revolutionary changes in the tree breeding due to the advanced biotechnological tools available world over. The further studies can be conducted to ascertain the biochemical or cellular changes responsible for flowering and fruiting in *Cordia myxa*.

## Conclusion

As a natural variant only seedlings of *Cordia myxa* produced functional flowers and fruits respectively at six months of age. This is the first report on the fruiting in the *Cordia myxa as* usual the flowering and fruiting takes place after many years of germination and on completion of juvenile stage and present specimen is extremely rare in trees species. However, it could be very useful for breeding and early evaluation of trees regarding floral biology and fruiting behavior. This is a rare plant on which studies with regard to fruiting can be conducted to ascertain the biochemical or cellular changes responsible for flowering and fruiting in *Cordia myxa*. Development of self breeding lines can be a key step in crop improvement programmes because can help in the production of true breeding varieties and reproducible hybrids in shortest possible time.

## Acknowledgements

The author is highly thankful to the department of sciences and technology GOI and H.P. Govt for providing financial help in conducting the research on the above stated topic.

## REFERENCES

Bernier, Georges Kinet, Jean-Marie, Jacqmard, Annie, Havelange Andree and Bodson, Monique. 1977. Cytokinin as a Possible Component of the Floral Stimulus in Sinapis alba Received for publication Plant Physiol. 60, 282–285

Bhatnagar, Prerak, Jitendra Singh, LK Dashora, MC Jain and CB Meena. 2015. “Genetic Variability Studies of lasoda (*Cordia myxa*) Genotypes for Jhalawar district in Rajasthan, India”. EC Agriculture 2.2 (2015): 300–306.

Cecich, R. A. 1981 “Applied Gibberellin Aq7 Increases Ovu-late Strobili Production in Accelerated Growth Jack Pine Seedlings,” Canadian Journal of Forest Research, vol. 11, no. 3, 1981, pp. 580–585.

Cecich, R. A. Kang, H. and Chalupka, W. 1994. “Regulation of Early Flowering in *Pinus banksiana*,” Tree Physiology, vol. 14, 275–284.

Furr, J. R., Cooper, W. C. and Reece, P. C. 1947. An Investigation of Flower Formation in Adult and Juvenile Citrus Trees. American Journal of Botany vol. 34, no. 1 (Jan., 1947), pp. 1–8.

Guo, J., Tian, L., Sun, X. Z., and Al, E. 2015. Relationship between endogenous polyamines and floral bud differentiation in *Chrysanthemum morifolium* under short-day conditions. Wonye kwahak kisulchi 33, 31–38.

Hilton. Brian, 2008. Applying Calcium Carbide to Induce Flowering in Pineapple. ECHO Development Notes (EDN) | EDN Issue :101.23–26.

Johnson, L. C. and Critchfield, W. B. 1978. “The Production of Functional Pollen and Ovules by Pine Seedlings Less than l-Year-Old,” Forest Science, vol. 24, 467–468.

Kumar, Rahul, Khurana, Ashima, Sharma, Arun K. 2014. Role of plant hormones and their interplay in development and ripening of fleshy fruits. Journal of Experimental Botany, 65(16) 4561–4575.

Lang A. 1965. Physiology of flower formation. In W. Ruhland, ed, Encyclopedia of Plant Physiology Vol 15. Springer-Verlag, Berlin pp 1380–1536.

Manisha J. Oza and Yogesh, A. Kulkarni. 2016. Traditional uses, phytochemistry and pharmacology of the medicinal species of the genus Cordia (Boraginaceae). Journal of Pharmacy and Pharmacology, 69, 755–789.

Peter, KV. 2007. “Horticulture Science Series”. Fruit Crops, New India Publishing Agency 3 : 353.

Pharis, R. P. and King, R. W. 1985. “Gibberellins and Repro-ductive Development in Seed Plants,” Annual Review of Plant Physiology, vol. 36, pp.517–568.

Sharma, Kamal, Thakur, Sanjeev, Sharma, Seema and Sharma, Som Dutt. 2012. A New Record on Flowering in Harar (*Terminalia chebula* Retz.) Seedling. American Journal of Plant Sciences, (3) 693–695.

Shwaish, Tarik and Al-Imarah, JM Faris. 2017. Chemical composition of *Cordia myxa* fruit: phytochemical screening and identification of some bioactive compounds. Int. J. Adv. Res. 5(9), 1255–1260.

Teich, A. H. and Holst, M. J. 1969. “Genetic Control of Cone-Clusters and Precocious Flowering in *Pinus sylvestris*,” Canadian Journal of Botany, vol. 47, no. 7, 1081–1084.

Tiren, L. 1935. Om granens kottsattning, dess periodicitet och samband med temperatur och nederbord. (On the fruit setting of spruce, its periodicity and relation to temperature and precipitation.) - Medd. Statens Skogsforsoksanst. 28: 413–524.

Tiren, L. 1935. Om granens kottsattning, dess periodicitet och samband med temperatur och nederbord. (Flowering and seed setting of spruce, its periodicity and relation to temperature and precipitation.) (In Swedish with English summary.) - Medd. Statens Skogsforsoksanstalt 28: 413–518.

Troup, R. S. 1921. “The Silviculture of Indian Trees,” International Book Distributors, Dehra Dun, pp-659.

Vakshasya, R. K. and Rawat, M. S. 1986. “Mutation for Precocious Flowering in *Dalbergia sissoo*,” Silvae Genetica, vol. 35, 247–248.

Xu, L. 2015. The effect of polyamineon flower bud differentiation and bud germination of chrysanthemum. Shandong Agric. Univ. 31–36.

Zhang, H. D. E., Harry, C. Ma, Hsu, C. Yuceer, C. Yu., Vikram V. Shevchenko, O., Etherington, E. and Strauss, S. H. 2010. “Precocious Flowering in Trees: The FLOWERING LO-CUST Gene as a Research and Breeding Tool in *Populus*,” Journal of Experimental Botany, vol. 61, (10), 2549–2560.

